# Out of thin air: surveying tropical bat roosts through air sampling of eDNA

**DOI:** 10.1101/2022.11.07.515479

**Authors:** Nina R. Garrett, Jonathan Watkins, Charles Francis, Nancy B. Simmons, Natalia V. Ivanova, Amanda Naaum, Andrew Briscoe, Rosie Drinkwater, Elizabeth L. Clare

**Author notes:** Corresponding Author: Nina R. Garrett^1^, 4700 Keele St., Toronto, ON, M3J 1P3, Canada.

## Abstract

Understanding roosting behaviour is essential to bat conservation and biomonitoring, often providing the most accurate methods of assessing population size and health. However, roosts can be challenging to survey. Roosts can be physically impossible to access or present risks for researchers and disturbance during monitoring can disrupt natural bat behaviour and present material risks to the population e.g. disrupting hibernation cycles.

One solution to this is the use of non-invasive monitoring approaches. Environmental (e)DNA has proven especially effective at detecting rare and elusive species particularly in hard-to-reach locations. It has recently been demonstrated that eDNA is carried in air and, when collected in semi-confined spaces can provide remarkably accurate profiles of biodiversity, even in complex tropical communities.

In this study we deploy novel airborne eDNA collection for air for the first time in a natural setting and use this approach to survey difficult to access potential roosts in the neotropics. Using airborne eDNA we confirmed the presence of bats in 9 out of 12 roosts. The identified species matched previous historical records of roost use obtained from photographic and live capture methods demonstrating the utility of this approach. We also detected the presence of the white-winged vampire bat (*Diaemus youngi*) which had never been confirmed in the area but was long suspected. In addition to the bats, we also detected several non-bat vertebrates, including the big-eared climbing rat (*Ototylomys phyllotis*), which has previously been observed in and around bat roosts. We also detected eDNA from other local species known to be in the vicinity. Using airborne eDNA to detect new roosts and monitor known populations, particularly when species turnover is rapid, could maximize efficiency for surveyors while minimizing disturbance to the animals. This study presents the first applied use of airborne eDNA collection for ecological analysis and demonstrates a clear utility for this technology in the wild.

## Introduction

### Bats and Their Roosts

Bat species are characterized by a wide variety of roosting ecologies (Fenton & Ratcliffe, 2010; Voss et al., 2016) utilizing caves, trees, man-made structures, cracks in rocks (Altringham, 2011), leaf litter (Mormann & Robbins, 2007), and even pitcher plants (Grafe et al., 2011). Some species modify the environment to create their roosts (e.g., creating leaf tents (Kunz, 1982) or excavating roosts within termite mounds (Esquivel et al., 2020)) and multiple species may use the same roost (Villalobos-Chaves et al., 2016; Kelm, Toelch & Jones, 2021). Bats require safe roosts that provide protection from predators with appropriate environmental conditions related to temperature and humidity. Bats may use different roosts at night or during the day at different times of year, for breeding, migration or hibernation (Altringham, 2011). Roosts are additionally important for mating and raising young, playing a key role in social interactions and maintaining populations (Humphrey, 1975; Kunz, 1982). Many bat species live in fission-fusion societies where roost switching is common (Patriquin & Ratcliffe, 2016) and supports larger social networks (Patriquin & Ratcliffe, 2016) but generates high individual turnover rates in roost occupancy (Aguirre, Lenstra & Matthysen, 2003; Patriquin & Ratcliffe, 2016) creating a challenge for conservation monitoring.

Understanding roosting ecology is important for bat conservation, especially as roosts are thought to be a limiting resource for some species (Humphrey, 1975; Aguirre, Lenstra & Matthysen, 2003; Voss et al., 2016). Roost surveys can inform decision making regarding the protection of bat habitat and roost loss prevention (Villalobos-Chaves et al., 2016), and can help understand and monitor community composition (Voss et al., 2016; Kelm, Toelch & Jones, 2021). Monitoring roost occupancy is one of the most effective ways to estimate the population sizes of some species (Kunz, 2003) and has been key to tracking the impact of disease dynamics e.g. white nose syndrome in North American populations (Janicki et al., 2015). Annual roost surveys are conducted in many regions to estimate population health (Kaarakka, 2020; Bat Conservation Trust, 2021) and in temperate zones, roost monitoring provides insight into migration stopovers and hibernating species (Klüg-Baerwald et al., 2017).

### Airborne eDNA Sampling for Roost Surveys

Traditional methods of roost surveying are challenging, expensive in time and cost, and can be invasive to the animals. Some roosts are physically inaccessible while others are too dangerous or toxic for humans to explore. It can be challenging to accurately determine species composition using existing methods (Behrens et al., 2017), and conventional methods additionally risk disturbing species which is especially detrimental to hibernating bats causing arousal and unnecessary use of fat reserves (Speakman, Webb & Racey, 1991). A non-invasive sampling method that does not require physical access to the bats could help overcome these challenges.

One way to increase roost monitoring efficiency is the use of environmental (e)DNA. eDNA is any genetic material not collected directly from the individual, and it has become a powerful tool in detecting organisms without physical access to individuals. Sampling eDNA from water or soil has become widespread (Thomsen & Willerslev, 2015) and collecting aquatic eDNA it is now a standard tool in monitoring aquatic ecosystems (Rees et al., 2014; Ruppert, Kline & Rahman, 2019). More unconventional methods interrestrial zones have targeted eDNA from spider webs (Gregorič et al., 2022) and snow tracks (Kinoshita et al., 2019) to learn about local ecology. Cavity roosts of bats have been suggested as an ideal target for terrestrial eDNA collections (Clare et al. 2021). The very reason that they are used by roosting bats – being enclosed and protected – may contribute to the longer-term preservation of environmental DNA which might otherwise degrade or be washed away (Mena et al., 2021) or become too dispersed to capture. Soil from caves has been used to detect cave-dwelling species, both those that are currently present and those from the recent past (Hofreiter et al., 2003), suggesting the presence of accumulating eDNA in these habitats.

Collecting and analysing airborne eDNA has been proposed as a method to monitor terrestrial animals (Barnes & Turner, 2015; Ruppert, Kline & Rahman, 2019). The first paper to demonstrate this technique targeted naked mole rats in artificial burrows (Clare et al. 2021) because of the perceived potential for eDNA to build up in an enclosed space. Airborne eDNA detection of vertebrates, insects and general biodiversity is in its infancy, but has already proven useful in detecting plant species missed using conventional sampling (Johnson et al., 2021). Airborne eDNA does not require access to the individual animal, reducing the risks associated with disrupting roosting bats, potentially allowing extended sampling times in otherwise inhospitable roosts, and potentially permitting sampling in roosts that are inaccessible using existing methods. The use of airborne eDNA to detect terrestrial vertebrates has been validated both inside and outside artificial dens in zoos (Clare et al., 2022; Lynggaard et al., 2022). Passive airborne dust collection methods sampling for weeks at a time have also been able to detect recent mammal activity in natural landscapes (Johnson et al., 2022). More recently Garrett et al. (2022), demonstrated that new prototype air sampling devices effectively detected eDNA from a diverse assemblage of Neotropical bats in an enclosed environment with remarkable accuracy. These findings validated the use of airborne eDNA in complex communities and suggest an effective novel approach for surveying roosts.

### Validating airborne eDNA for small cavity roost surveys in the neotropics

Given its potential demonstrated through previous pilot studies (Clare et al., 2021; Serraro et al. 2021, Clare et al., 2022; Lynggaard et al., 2022, Garrett et al. 2022), our objective was to evaluate airborne eDNA as an applied survey tool for a set of neotropical bat roosts in the first targeted deployment of airborne eDNA sampling in a truly natural setting. Our goals were to assess eDNA as a roost survey method and to develop a profile of roost use in Central America. Neotropical bat roosting behaviour is complex and understudied (Fenton et al., 2001; Villalobos-Chaves et al., 2016), and monitoring using airborne eDNA could be a game-changing approach to this field. The bat fauna in our study site has been well documented for over a decade using live capture methods (i.e., mist nets, hand nets, and harp traps; Fenton et al., 2001; Herrera et al., 2018)) and camera traps (Rydell et al., 2022) giving us *a priori* knowledge of the local bat fauna as well as roosting ecology of many species. This creates an ideal study system in which to test the application of airborne eDNA during roost surveys and directly compare to the known species inventories. Using this system, we collected airborne eDNA from a variety of natural and man-made roosts. We test the hypothesis that airborne eDNA collects in sufficient quantities in natural roosts to document roosting ecology of cavity-roosting neotropical bat species.

## Materials & Methods

### Study Site

This study was conducted in late April and early May 2022 in and around the riverine forest of Lamanai Archaeological Reserve (LAM) and the nearby forest fragment of the Ka’kabish Archeological Research Project (KKB) in the Orange Walk District of Belize. Both sites are ancient Maya cities that have become overgrown with semi-deciduous tropical forest. LAM contains excavated ruins that are open to the public and preserves approximately 450-ha of tropical forest adjacent to the freshwater New River lagoon (Herrera et al., 2018). The forest fragment at KKB is substantially smaller (45-ha) and is entirely surrounded by agricultural land (Herrera et al., 2018); it is not open to the public. Both LAM and KKB are surrounded by a matrix of agricultural fields, pastures, farms, and villages. Work in this area was conducted under Belize Forest Department permits FD/WL/1/21(12) and FD/WL/1/21(18), and Belize Institute of Archaeology permit IA/H/1/22(03).

### Roost Surveys

We sampled twelve known or suspected bat roosts in LAM, KKB, and nearby local farms and villages. These consisted of four tunnels carved into Maya ruins at KKB by archaeologists and looters (Fig. 1B, 1C); one large cistern in LAM; one attic in a house in Indian Church Village; four hollows in large trees in the LAM; one natural cave of uncertain size in secondary forest (Indian Creek Cave) and one relatively small natural cave in a small cleared hill in a pasture (Schoolhouse Cave ; Fig. 1E), both in the vicinity of Indian Creek (Table 1). At the Schoolhouse Cave, we placed our samplers 3 to 5 metres inside and spanning the width of the cave (3 to 4 metres) (Fig. 1E). Indian Creek Cave had a steep vertical drop at the entrance and the White Room roost was covered by an unstable tin roof, so for safety reasons we did not enter these roosts. In both cases we placed our samplers near the entrance. The Red Room roost (Fig. 1A) was approximately 4 to 5 meters in height and 2 to 3 meters in width. The White Room roost (Fig. 1B) was part of the same ruin as the Red Room roost, but on the opposite side of the structure. Two of the artificial tunnels in KKB, Plaza Tunnel, and *Natalus* Tunnel, were accessible but were relatively narrow, about 1 to 1.5 metres across (Fig. 1C) and 1.5 to 2 meters tall. Three of the four hollow trees were large Guanacaste trees around LAM (Fig. 1D, 1F), with accessible openings large enough to set up samplers inside the hollows. The Museum Tree had a much smaller opening, only a few centimetres wide and was located near the LAM museum. For this roost, the sampler was positioned facing inwards just outside the entrance slot.

**Table 1.** Sampling effort at each of the roost sites, indicating number of 12V air samplers run at one time, number of separate days they were deployed, and the runtime of each.

| Type | Roost | # Samplers | # Deployments | Time Sampled (hrs) |
| --- | --- | --- | --- | --- |
| Natural Caves | Indian Creek Cave | 2 | 1 | 8 |
|  | Schoolhouse Cave | 6 | 2 | ~24 |
| Artificial Tunnels (KKB) | Red Room | 1 | 1 | ~24 |
|  | Plaza Tunnel | 1 | 2 | ~24 |
|  | *Natalus* Tunnel | 1 | 2 | ~24 |
|  | White Room | 1 | 1 | ~24 |
| Other Human-made | Helen's House | 1 | 1 | ~24 |
|  | Cistern (LAM) | 2 | 1 | 6 |
| Tree Roosts (LAM) | High Temple Hollow Tree | 1 | 2 | 6 |
|  | Museum Tree | 1 | 1 | 6 |
|  | Sugar Mill High Tree | 1 | 3 | 6 |
|  | Sugar Mill Low Tree | 1 | 2 | 6 |

**Figure 1.**
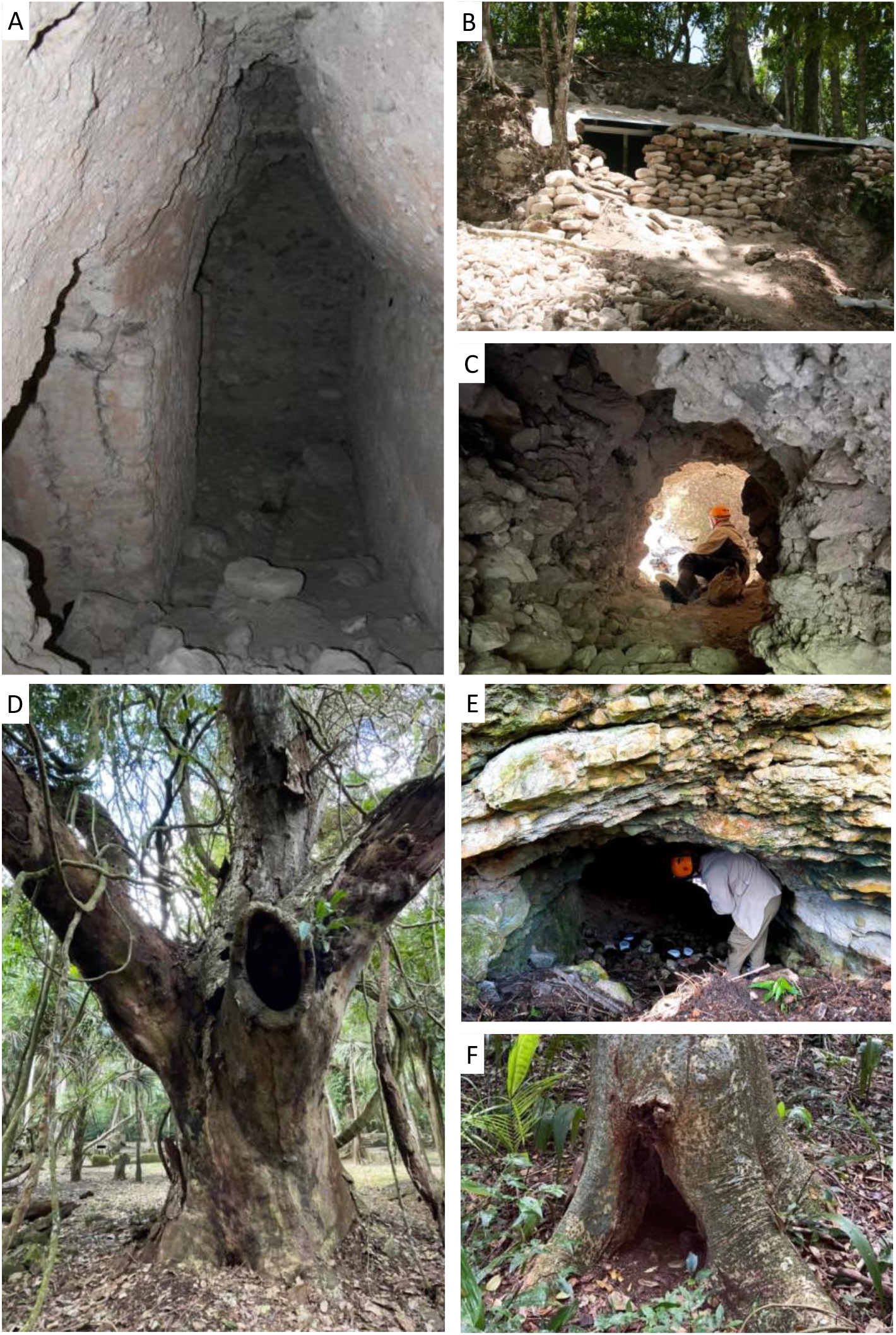
Twelve natural and manmade roosts were surveyed using airborne eDNA. These included manmade looters tunnels in Maya ruins (A, C), archaeological excavations (B), tree roosts (D, F), and natural caves (E). Samplers were deployed inside roosts (e.g., E) and left for up to 24 hours to filter air (Images by Helen Haines (B), Elizabeth Clare (A,E,F), and Nancy Simmons (C,D)).

We filtered roost air by deploying 12vLarge samplers as described by Garrett et al. (2022) in each roost (Table 1, Fig. 1). We cleaned samplers with a 50% bleach solution and then water before each use, and KN95 masks and gloves were worn by researchers when handling filters. Sampling varied across roost sites based on roost size, access, and weather. For example, samplers in the LAM could only be deployed when the Reserve was closed, between late afternoon and dawn, limiting sampling hours at roosts in that area. Samplers also could not be left out during heavy rain in LAM as they were uncovered. In total we ran samplers for approximately 24 hours in six of the roosts, for six hours in five roosts, and for eight hours in one roost (Table 1). In Schoolhouse Cave, the cave was sampled for approximately 24 hours starting at about 9:30 in the morning; however, the filters were changed in the late afternoon, producing one set of daytime and one set of overnight samples. This resulted in 27 samples collected in total across 12 roost sites. Filters were removed from samplers, folded so the exposed “disk” collection surface was on the inside, and placed in sterile bags before they were frozen for storage.

### DNA Extraction

Sample processing was performed in decontaminated, UV sterilized Biosafety cabinet to limit sources of extraneous eDNA. The decontamination protocol involved cleaning the surfaces of the Biosafety cabinet (including the space under the grill) with 1% Virkon, followed by 70% Ethanol. A head cover, mask, lab coat, gloves and sleeve covers were used to minimize human DNA load during subsampling and DNA extraction. Prior to extraction we unfolded the filters (one at a time) and cut out a half circle from the centre of each filter disk using sterile scissors. We then cut each half circle into segments and placed these in a 5mL Eppendorf tube with 4mL of PBS. We soaked these overnight while incubating them at 56°C in using a rotary wheel hybridization oven. Following incubation, we transferred 1000μL of the PBS solution from each filter sample to a 1.5mL DNA LoBind Eppendorf tube and spun it at 6000 (xG) for three minutes. We pipetted the liquid into an empty 5mL DNA LoBind Eppendorf tube, leaving any precipitate behind. We repeated this process until all the PBS was removed from the first 5mL DNA LoBind Eppendorf tube and the precipitate was concentrated into one Eppendorf tube. For all subsequent steps we treated the precipitate as the “tissue” and DNA was extracted using a Qiagen Blood and Tissue Kit (Qiagen) following manufacturers guidelines with the exception of the elution step, where we incubated the buffer at 56°C and eluted the DNA in 100μL of elution buffer. We processed extraction blanks using only the solutions in the kit as an extraction negative control. We froze the extracted DNA prior to PCR.

### PCR and Sequencing

PCR reagent preparation was done in the AirClean PCR cabinet located in the ISO 7 Clean Room at NatureMetrics laboratory in Guelph, Ontario. Head covers, lab coat, gloves, sleeve covers, and boot covers were worn in a Clean Room to minimize human DNA load. PCR protocols follow those described by Garrett et al. (2022). PCR setup (adding DNA to plates prepared in ISO 7 clean room) was performed in the PCR-free room in the AirClean PCR cabinet decontaminated and UV sterilized as described above, using the same PPE with the exception for boot covers. We preformed three technical replicates PCRs using the mam1 and mam2 primers (∼90bps) (Taylor, 1996; Calvignac-Spencer et al., 2013) modified with Illumina adaptors. We included negative (no template) and positive (*Pteronotus psilotis)* controls and visualized all PCR products, including all controls (positive, negative, and extraction blank) using an Invitrogen E-Gel™ 96 Agarose Gels with SYBR™ Safe DNA Gel Stain, 2%, run for eight minutes on the E-Gel™ Power Snap Plus Electrophoresis System. All PCR products were sequenced on the Illumina MiSeq by the NatureMetrics laboratory in Guelph, Ontario using the sequencing protocols of Garrett et al. (2022). Reads were demultiplexed and exported as FASTQ files in preparation for bioinformatic analysis.

### Bioinformatic Methods

During standard bioinformatic processing of the data, we identified several ASVs (amplicon sequence variants) which showed evidence of unexpected primer combinations. The PCR products had been sequenced with an unrelated data set by pooling amplicons from different areas of the genome for barcoding with the same Illumina tag. This resulted in a small number of sequences with a forward primer of one amplicon and a reverse primer of the other, which made it impossible to automate primer removal. To correct this, we processed the data using the DADA2 pipeline as described by Garrett et al. (2022) but without primer removal to generate ASVs with primers still attached. We then examined these ASVs manually in BIOEDIT (Hall, 1999) and separated the 16S reads from non-16S (the unrelated project which shared the sequencing run) reads based on known nucleotide signatures of the two regions amplified which are quite distinct. We identified a small number of ASVs which had this mixed or incomplete primers, likely from primer leftover during library building when independent projects were pooled for barcode addition. These ASVs represented less than 0.006% of the total data and they were discarded. We identified and removed the intact primers from the remaining 16S ASVs manually in BIOEDIT.

We compared the trimmed ASVs to the full NCBI nucleotide collection using BLAST. We removed all ASVs matched to human DNA and, based on full negative control filtering (Garett et al. 2022), we discarded all ASVs with read counts lower than 21, the highest read count identified in any negative control replicate after removal of human DNA. These 21 reads were identified as *Pteronotus psilotus* the species we used as our positive control. All ASVs greater than 96% identity (100% overlap) were retained for further examination. We also retained ASVs matched to the bat *Chrotopterus auritus* and primate *Alouatta spp*. based on lower percentage matches. *Chrotopterus auritus* is considered to be an unresolved cryptic species complex with at least three distinct mitochondrial lineages with as much as 16% sequence divergence between Central and South American lineages (Clare et al., 2011). The closest match to reference material on NCBI comes from a specimen from Peru (AMNH Mammalogy 280554) and thus a more relaxed match of 93-94% with no other similar reference was retained. Several ASVs match to *Alouatta palliata* at 92.5%. This species is not found in Belize, but the related *Alouatta pigra* is common in our research area. The taxonomy of *Alouatta* is complex and has recently undergone revisions (Doyle et al., 2021). It is not clear if any *Alouatta pigra* 16S references are contained in the Genbank nucleotide collection (the name does not exist in Genbank). We retain *A. pigra* for these reads as the mostly likely identification. Six of the samples (two each from Sugar Mill High Tree, Sugar Mill Low Tree, and Schoolhouse Cave) had been sequenced previously (Garrett et al. 2022) and were included at the analysis stage.

### Sample Coverage and Day vs. Night Detections

We could easily enter the Schoolhouse Cave and the floor area of the roost was large, allowing for a greater sampling effort. Therefore, from this roost we estimated the effect of sampling effort on taxonomic recovery (Fig. 1E). Using a Hill number approach, we generated diversity accumulation curves at three different diversity orders of (q). These Hill numbers are equivalent to the commonly used diversity indices: species richness (q=0), the Shannon index (q=1), and the Simpson index (q=2). The diversity profiles were generated with 95% confidence intervals using the iNEXT package (Chao et al., 2014; Hsieh, Ma & Chao, 2022) in RStudio (RStudio Team, 2021). Following the protocol described by Chao et al. (2014), curves were extrapolated to double the size of the observed value. With limited resources and access to multiple sites, we also ran a test of the difference in detections over one night of sampling in the Schoolhouse Cave. We performed a paired t-test on the mean number of species detected to determine if more bat species were detected at night than during the day. We also compared whether more non-bat vertebrates were detected during the day then at night. We tested the homogeneity of the data using the Bartlett test and the distribution using the Shapiro-wilks test.

## Results

### Species Detections

Of the 207 ASVs identified after bioinformatic processing, we retained 138 after filtering and removal of the positive control. We identified these as coming from 25 taxa, including 12 bat taxa, four amphibian species, three non-bat native mammal taxa, and six domestic mammals (Table 2). One bat taxon (*Molossus)* could only be identified to the genus level as two local species have very similar DNA sequences (although a photograph at the roost suggests it may have been *M. alverezi* (Fig. 2H; see discussion).

**Table 2.** Summary of roost survey detections grouped by roost type. Taxa with a % match in GenBank (NCBI) lower than 95% are denoted with a *.

| Type | Roost | Bats | Other Vertebrates |
| --- | --- | --- | --- |
| Natural Caves | Indian Creek Cave | *Carollia perspicillata*<br>*Desmodus rotundus*<br>*Glossophaga mutica*<br>*Natalus mexicanus* | *Bos taurus* (cow) |
|  | Schoolhouse Cave | *Carollia perspicillata*<br>*Glossophaga mutica*<br>*Natalus mexicanus*<br>*Saccopteryx bilineata*<br>*Sturnira parvidens*<br>*Trachops cirrhosus* | *Alouatta pigra* * (Yucatan black howler monkey)<br>*Bos taurus* (cow)<br>*Canis spp.* (canine spp.)<br>*Equus caballus* (horse)<br>*Otodylomys phyllotis* (big eared climbing rat)<br>*Leptodactylus fragilis* (white-lipped frog)<br>*Ovis aries* (sheep)<br>*Scinax staufferi* (Stauffer's tree frog)<br>*Sus scrofa* (pig)<br>*Sylvilagus floridanus* (eastern cottontail)<br>*Trachycephalus typhonius* (veined tree frog) |
| Artificial tunnels (KKB) | Red Room | No detections | *Canis spp.* (canine spp.) |
|  | Plaza Tunnel | *Carollia perspicillata*<br>*Saccopteryx bilineata* | *Otodylomys phyllotis* (big eared climbing rat)<br>*Sus scrofa* (pig)<br>*Trachycephalus typhonius* (veined tree frog) |
|  | Natalus Tunnel | *Carollia perspicillata*<br>*Glossophaga mutica*<br>*Natalus mexicanus*<br>*Sturnira parvidens*<br>*Trachops cirrhosus* | *Bos taurus* (cow)<br>*Canis spp.* (canine spp.)<br>*Dendropsophus microcephalus* (yellow tree frog)<br>*Otodylomys phyllotis* (big eared climbing rat)<br>*Ovis aries* (sheep)<br>*Leptodactylus fragilis* (white-lipped frog)<br>*Sus scrofa* (pig) |
|  | White Room | *Chrotopterus auritus* * | *Bos taurus* (cow)<br>*Otodylomys phyllotis* (big eared climbing rat) |
| Other Human-made | Helen's House | *Glossophaga mutica* | *Bos taurus* (cow)<br>*Canis spp.* (canine spp.) |
|  | Cistern (LAM) | No detections | *Bos taurus* (cow)<br>*Canis spp.* (canine spp.) |
| Tree Roosts (LAM) | High Temple Hollow Tree | No detections | No detections |
|  | Museum Tree | *Molossus spp.* | *Alouatta pigra* * (Yucatan black howler monkey) |
|  | Sugar Mill High Tree | *Desmodus rotundus*<br>*Diaemus youngi*<br>*Saccopteryx bilineata* | *Bos taurus* (cow)<br>*Otodylomys phyllotis* (big eared climbing rat)<br>*Ovis aries* (sheep) |
|  | Sugar Mill Low Tree | *Pteronotus fulvus*<br>*Sturnira parvidens* | No detections |

**Figure 2.**
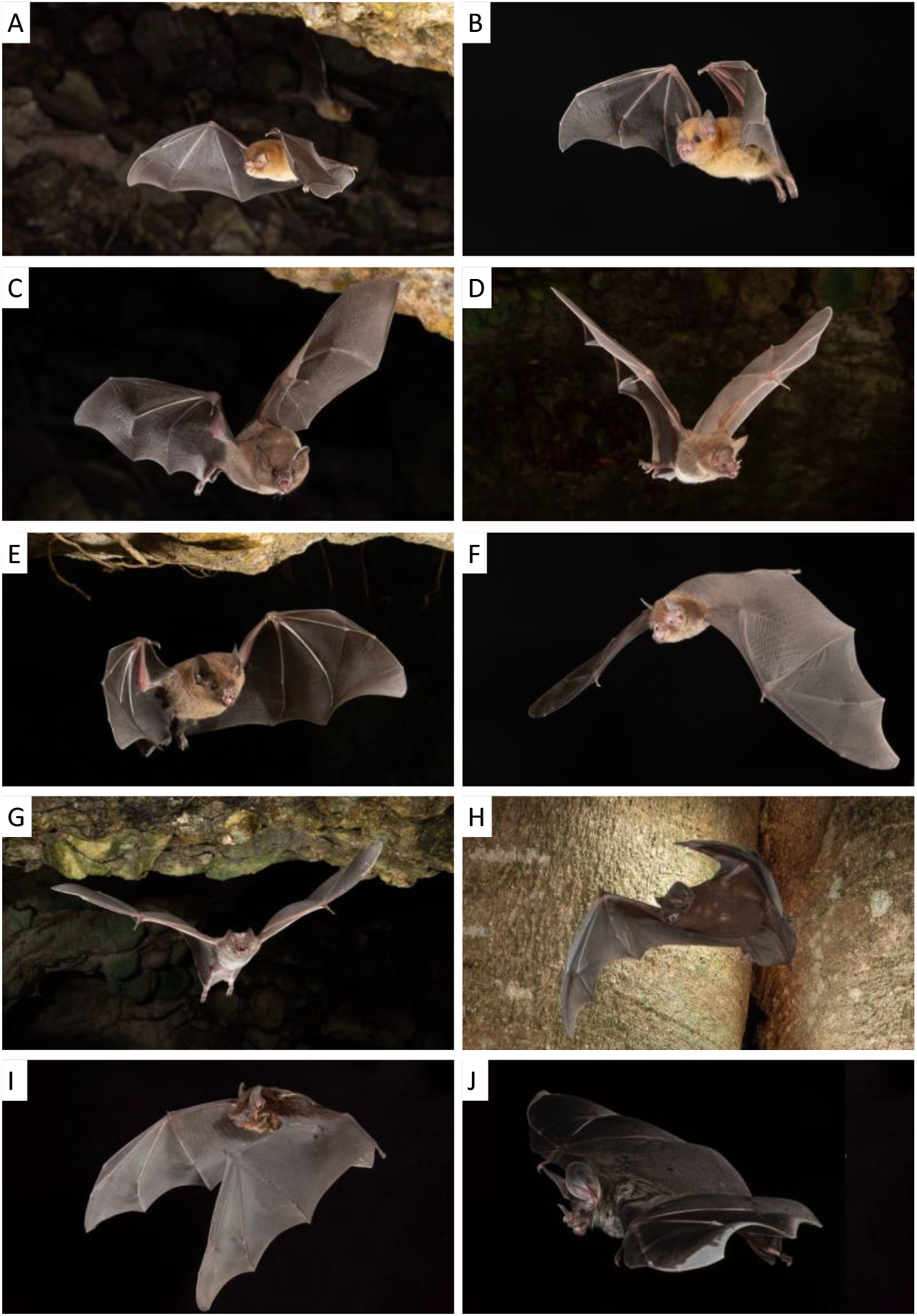
With the exception of *Diaemus youngi*, all bats detected using airborne eDNA have also been documented in the study area from camera traps at roost exits, from mist net captures and/or from being captured in roosts. *Natalus mexicanus* (A), *Glossophaga mutica* (C), *Carollia sp*. (E), *Desmodus rotundus* (G) and *Molossus* cf. *alverezi* (H), were detected using camera traps exiting at least one of the roosts where their DNA was detected. *Trachops cirrhosis* (D) was detected with a camera trap at a different cave roost. *Pteronotus fulvus* (F), *Sturnira parvidens* (B) and *Saccopteryx bilineata* (I) were all captured regularly, while *Chrotopterus auritus* (J) has only been detected at an artificial tunnel we did not sample this year using traditional methods; photographs of these last 4 species were taken in a studio setting (images A-I by Charles M Francis, F by M. Brock Fenton.

The natural cave roosts recovered the highest overall richness of taxa from DNA with 17 species being identified. These were six bat species, three non-bat native mamals, five domestic mammals, and three amphibian species (Table 2). From a cave roost, we also retrieved the only detection of the frog species *Scinax staufferi*, in the Schoolhouse Cave (Table 2). In the tree roosts, we detected six bat species, two of which, *Diaemus youngi* (Sugar Mill High Tree) and *Molossus sp*. (Museum Tree), were not detected in any of the other roosts (Table 2). We also detected DNA from two non-bat native mammals and two domestic mammals in the tree roosts. We detected DNA from seven bat species, one non-bat native mammal, two amphibians, and four domestic mammals in the artificial tunnels. One of the frog species, *Dendropsophus microcephalus* (*Natalus* Tunnel), and one of the bat species, *Chrotopterus auritus* (White Room), were detected only in the artificial tunnels (Table 2). We detected DNA from one bat species and two domestic animals in the other man-made roosts. While DNA from domestic animals is almost certainly coming from the surrounding habitat, we can confirm the presence of the small mammals and bat species in the vicinity, and often in the same roosts, based on visual sightings, captures in nets nearby, and/or photographic records (Fig. 3).

**Figure 3.**
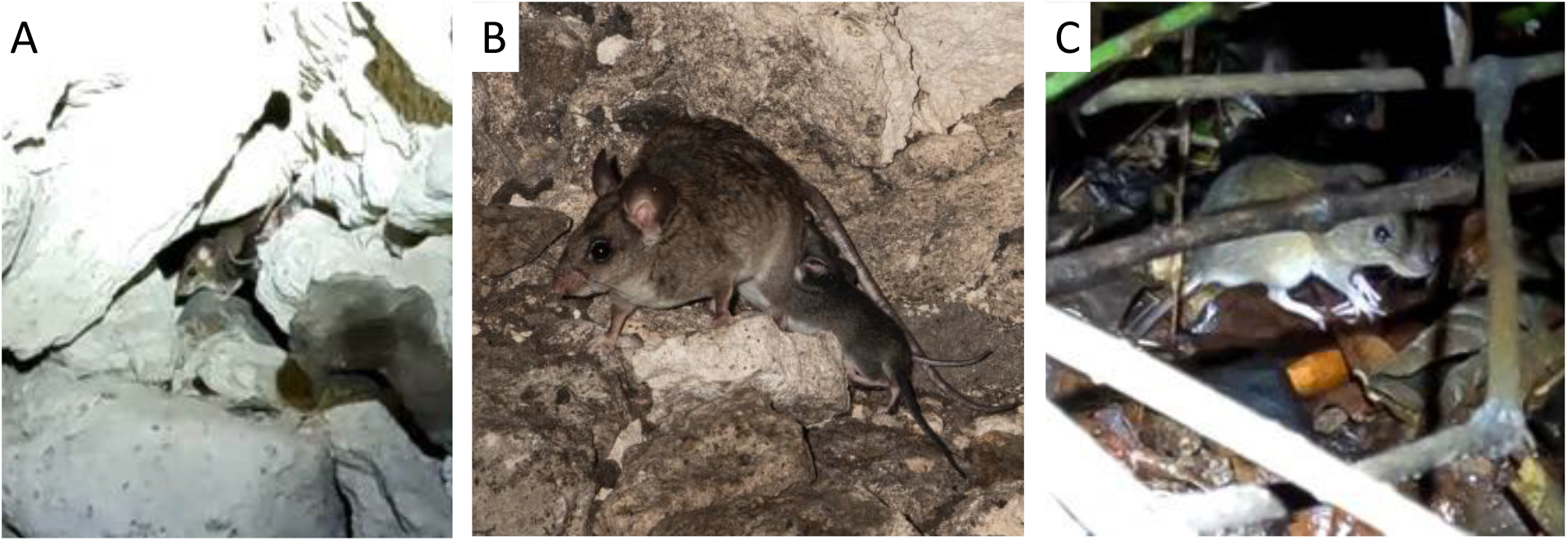
The big eared climbing rat *Ototylomys phyllotis* has been seen and photographed sharing roosting areas with bats in KKB (A, B) and in the area around the tree roosts of the LAM (C) in previous years and was both seen and recorded on acoustic equipment during the current field season. The DNA of *Ototylomys* was detected in both natural and man-made bat roosts and a hollow tree. This suggests widespread roost sharing behaviour for this species (images A&C Elizabeth L. Clare, B by Sherri Fenton).

### Sample Coverage and Day vs. Night Detections

The diversity accumulation curves show that within the Schoolhouse Cave sampling effort was sufficient to detect the majority of bat species in the cave; when we consider all orders of diversity (q=0,1,2), they all begin to plateau (Fig. 4). In contrast, when looking at the species richness accumulation (q = 0) for total diversity (bats and other vertebrate species) the curve does not yet reach an asymptote, indicating increased sampling may add more species. However, for the other orders of diversity (q=1,2) the curves plateau. This diversity profile indicates that most of the common species have been captured by the sampling effort, as q=1 can be considered the effective number of common species and q=2 the effective number of dominant species (Hsieh, Ma & Chao, 2016). But the mismatch with q=0, and with the bat-only profile could indicate that the current sampling was insufficient for rare non-bat vertebrates (Fig. 4). There was no statistically significant difference between the mean number of bats (t_5_= -1.19, p=0.14) or other vertebrate (t_5_=0.44, p=0.34) species detected during the day compared to eDNA accumulating during the night (Fig. 5). Both the normality and homogeneity assumptions were met. See appendix 1 (Fig. S1 and S2) for detections by sampler.

**Figure 4.**
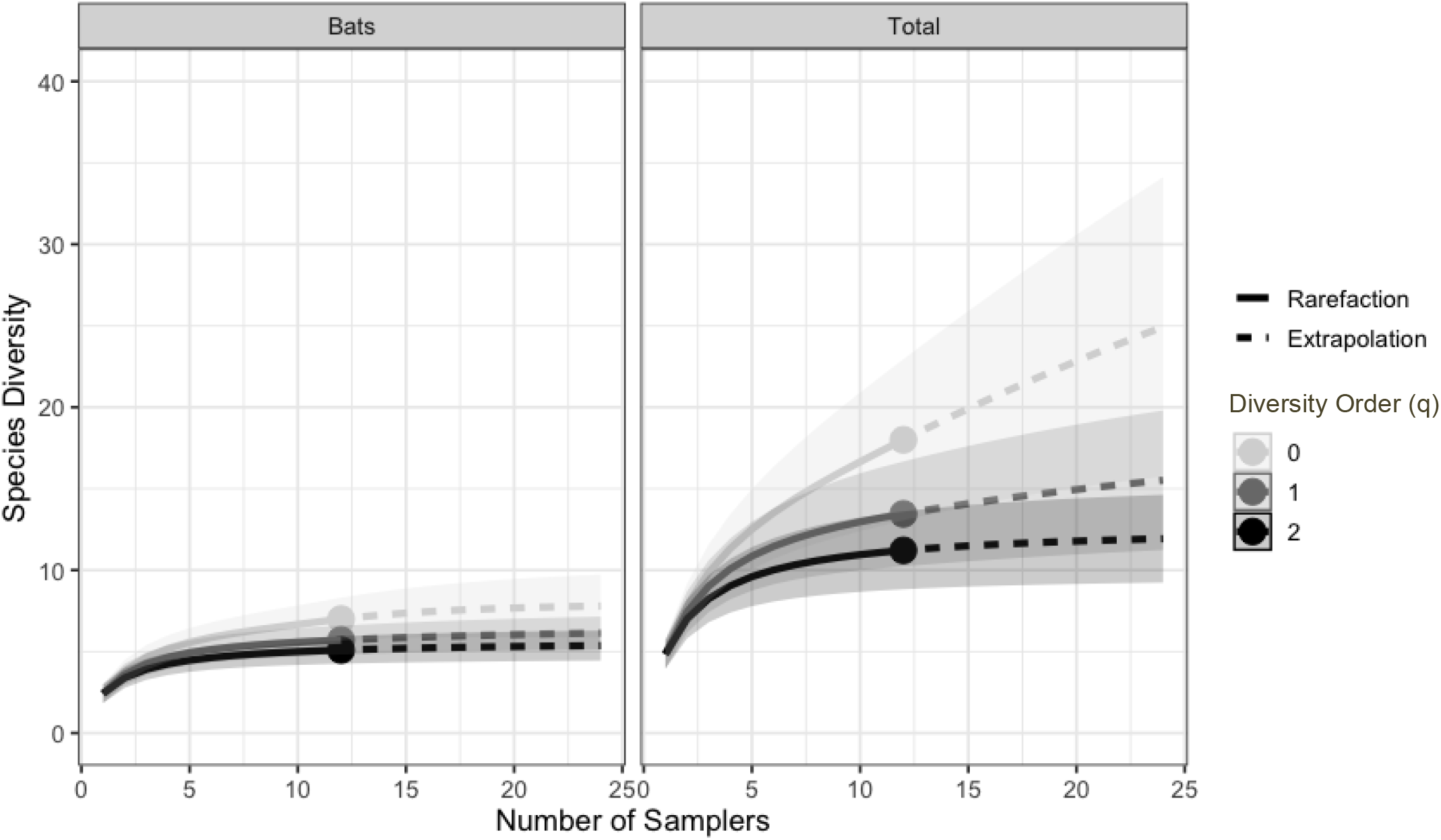
Accumulation curves for bat diversity and total diversity for three orders equivalent to species richness (q=0), the Shannon index (q=1) and the Simpson index (q=2). Including 95% confidence intervals and extrapolated to double the observed value (solid circle).

**Figure 5.**
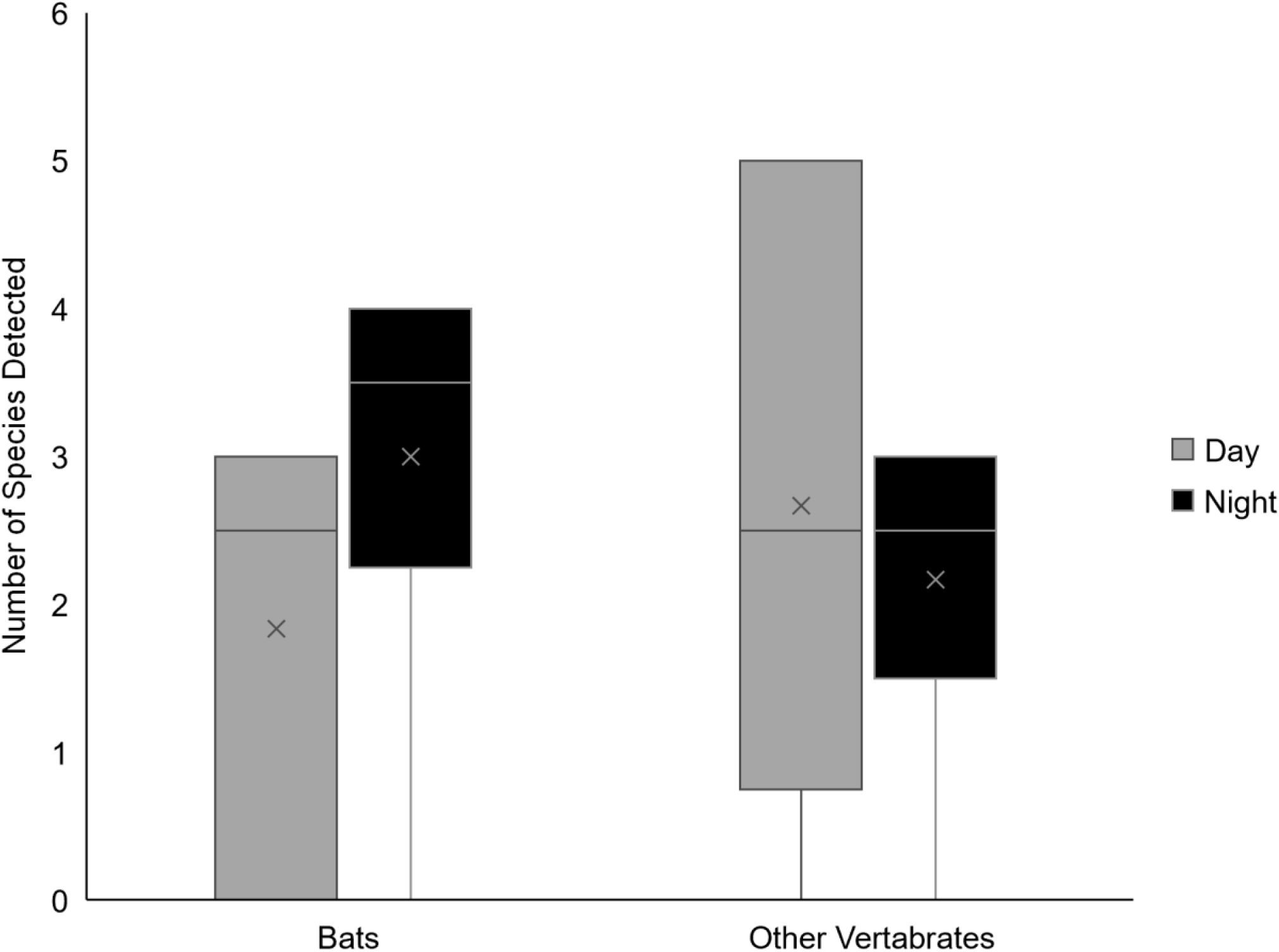
The mean (x) number of species detected during the day (grey, n=6) and night (black, n=6) for bat species (t_5_= -1.19, p=0.14) and other vertebrates (t_5_=0.44, p=0.34) at the Schoolhouse Cave.

## Discussion

In this study, we conducted the first real-world test of airborne eDNA sampling for applied ecological analysis of a wild terrestrial vertebrate community. Our goal was to document roosting ecology of neotropical bat species in small cavity roosts using non-invasive air sampling which minimizes any disturbance to the animals. In Belize, we sampled air from 12 potential roosts and were able to confirm bat occupancy in 9 of these, including at least one in each roost type (natural caves, artificial tunnels, other human-made structures, and tree cavities) therefore confirming that nine of the roosts were currently occupied. Of the three roosts without detections, one has never been surveyed (High Temple Hollow Tree) and so we had no evidence it was a roost, one is only known to be used as an occasional day roost (Cistern) and one (Red Room) had no bats present at any time we visited this year and no evidence of occupation (e.g., no guano, no acoustic recordings). Overall, we detected 25 taxa including bats, co-habiting mammals, and a selection of other known local animals, demonstrating that airborne eDNA can be used to detect bats in their roosts as well as other vertebrates in the surrounding area. These results showcase airborne eDNA’s potential to survey and monitor difficult to access locations with considerable efficiency.

### Roost Species Assemblages

We detected 12 bat taxa using airborne eDNA, several of which were documented at the same roosts using camera traps, and all but one of which have been captured in the vicinity. The one exception was *D. youngi*, the white-winged vampire, which was detected in a tree roost from DNA but has not previously been photographed or netted in the area, despite more than a decade of survey effort of at least 2 weeks per year. Despite no local records, the species range is believed to overlap this in area, and it was considered “likely to be present” with the local key to bats including a warning that it is suspected to be present (Clare & Simmons, 2021). Thus, its detection is a confirmation rather than surprise. This detection was also made independently in all three samplers at the location with a large read count providing robust evidence for its presence. Using airborne eDNA we also detected a potential new roosting type in the area for a well-known species *Sturnira parvidens* in the artificial tunnel and in the Schoolhouse Cave. Although regularly captured, this species had not previously been detected in cave-like roosts locally, it has only been documented in tree roosts (Fenton et al., 2000, under the name *S. lilium*). Whereas our DNA survey found evidence of this species in the *Natalus* Tunnel and in the Schoolhouse Cave.

The detection of rare or elusive species is a significant advantage of eDNA methods. Aquatic eDNA has been highly successful in detecting species in locations where they were not known to be present. For example, aquatic eDNA samples taken from caves in Croatia detected the presence of the IUCN red-listed amphibian, *Proteus anguinus*, for the first time in five different caves (Vörös et al., 2017). In Nova Scotia’s Kejimkujik National Park and Historic Site, aquatic eDNA detected the threatened Blanding’s turtle (*Emydoidea blandingii*) as well as two invasive species (chain pickerel (*Eesox niger*) and smallmouth bass (*Micropterus dolomieu*)) in locations where they were not known to occur (Loeza-Quintana et al., 2020). Our detection of *S. parvidens* and *D. youngi* in previously unknown roosting locations is a clear demonstration of the complimentary potential of this technique for assessing roosting ecology, even in locations which have been studied extensively using other methods.

We detected 12 non-bat vertebrates using airborne eDNA. Five of these were domestic animals whose DNA likely drifted into the roosts from the surrounding farmland. This drift is best showcased by the detection of cow DNA (*Bos taurus*) in the attic roost where it was clearly not possible for the animal to be physically present. Cows are abundant in the surrounding area, thus the detection of trace eDNA from such common species is likely going to represent a consistent false positive in many monitoring activities. In our recent validation of this sampling method for complex tropical bat communities (Garrett et al. 2022) we also detected commonly-known local species drifting on the wind which may suggest that, particularly for common species, pinpointing the source of an eDNA signal may be difficult. A similar problem is found in aquatic surveys where signals are occasionally found far from their source. In one study of the drift potential of DNA, Jane et al. (2015) were able to detect trout eDNA in streams at least 239.5m away from its source. Althought there is little research investigating the extent of such dynamics using airborne eDNA, our results indicate that such transport is likely for common speices. While most signals appeared quite localised, it will be difficult to trace all sources and more work on determining drift dynamics of airborne eDNA is required.

Of the remaining seven species detected in our study (four amphibians and three non-bat native mammals), it is likely that *Alouatta pigra* (the Yucatan black howler monkey) and *Sylvilagus floridanus* (easter cottontail) were also detected using DNA which drifted into the roosts. The four amphibian species and remaining mammal (*Ototylomys phyllotis*, big-eared climbing rat) may have been inside the roosts thus making their detections in the roosts true detections rather than caused by DNA drift alone. In particular, *O. phyllotis* has been previously observed and photographed in or around these roosts, and our data further confirms this roost-sharing behaviour in both tree and cave roosts (Fig 3). Non-target detections such as these suggest that airborne eDNA could be used not only to target one taxon, but document larger ecosystem -level community assemblages.

### Roosting Behaviour

Four of the bat species detected (*Carollia perspicillata, Desmodus rotundus, Glossophaga mutica, Trachops cirrhosis*) are known to use multiple roost types in the Neotropics in general (Reid, 2009) and at our study site in particular (Herrera et al., 2018) so their detection in multiple locations is not surprising. For example, *G. mutica* was detected in all the roost types except for the tree roosts (though we have previously caught them in Sugar Mill High Tree), and it was the only species detected in the attic roost at Helen’s House. *Natalus mexicanus* was only detected in cave or cave-like roosts (artificial tunnels). This behaviour is supported by other observations of these bats preferring to roost in these roost types (López-Wilchis, Torres-Flores & Arroyo-Cabrales, 2020). Despite photographic documentation and captures in hand nets in our study area, *Natalus* has never been caught in a mist net at this location in a decade of surveys. *Saccoperyx bilineata* was observed roosting at the entrances of the artificial tunnels and we have observed them roosting in the sugar mill structures near the tree roost where they were detected. They have been observed emerging from tree roosts in our study area and are known to roost in hollow trees elsewhere in the neotropics (Villalobos-Chaves et al., 2016; Voss et al., 2016), thus their detection in both the artificial tunnels where they were seen, and in the hollow tree roosts, is consistent with documented roosting behaviour. While it is believed that *S. parvidens* do sometimes roost in caves (or cave-like structures) as our data suggested and often co-roosts with other bats, they had not previously been observed doing this in the local area.

Historically, we have only found them in tree roosts. In contrast, *Pteronotus fulvus* is thought to prefer cave roosts (Willson & Mittermeier, 2019), but was only detected in a tree roost (Sugar Mill Low Tree) in our study. While we cannot confirm the detections the *S. parvidens* or *P. fulvus* with capture data from this year, the behaviour would not be surprising given the roosting patterns of congeners which also utilize tree roosts occasionally (Voss et al., 2016). However, it is also possible we are detecting eDNA moving from the local area into the roost, and more documentation of airborne eDNA movement patterns is required to determine the most likely explanation. We detected *Chrotopterus auritus* in only one roost, where it was the only species detected. This species, which is a large carnivorous bat, often roosts alone and while this roost was not surveyed by camera or netting during this field season because of safety concerns, the species has been caught in the same roost at that location in previous years (Brigham et al., 2018).

It should be noted that the *Molossus spp*. detection could not be identified to species. In a large survey (Clare et al. 2011) using DNA barcodes it was noted that while most central and South American molossids can be differentiated using mtDNA, the divergences are very minor. Given the small fragments amplified and sequenced in this study, we could not confidently identify the species. Perfect matches might be reliable, but more assessment using short reads is necessary. In this location the most reliable external character to differentiate the two regularly captured *Molossus* species is the white fur base in *Molossus alvarezi* and dark fur base in *Molossus nigricans* (Loureiro, Engstrom & Lim, 2020)). Based on camera trap images (e.g., Fig. 2H) we suspect that the *Molossus spp*. detected in the tree roost was *M. alverezi*. In that picture the fur has been parted by the air currents and a white base appears visible and more distinct than the pale skin under dark fur of *M. nigricans* would be.

### Airborne eDNA as a Roost Survey Tool

The use of airborne eDNA to study roosts, hollows, and burrows was cited as an ideal application in the first proof-of-concept of airborne eDNA detection of mammals (Clare et al. 2021). Our current study highlights the strong potential of this application with the first test of air-based bat roost surveys under natural field conditions. One of the most obvious advantages of this eDNA approach is that it enabled us to survey areas that were largely inaccessible and detect species that were not detected in our study area using other methods. The entrance to Indian Creek Cave drops steeply into the ground, making it difficult to enter the cave to survey bats. Similarly, it is not possible to enter the tree roosts (Fig. 1D and F) to visually identify species because the entrances and spaces used by the bats are, in many cases, too small to permit human entry. However, we were able to easily insert our small filter units into the entrances of these roosts and, using airborne eDNA, determine that these roosts were occupied and provide a basic list of the inhabitant species. Without this approach at least some of these roosts could not have been surveyed. For example, we were able to survey the tin roof structure (Fig. 1A) which was deemed too unstable to enter and thus unsafe to survey using nets or even photographic equipment. It also allowed us to detect *D. youngi*, a species not previously detected in the area but predicted to be present. The ability to detect elusive species is one of the main advocated benefits of eDNA. Aquatic eDNA studies have detected rare and elusive fish (Weltz et al., 2017; Nester et al., 2022), amphibians (Plante et al. 2021), birds (Neice & McRae, 2021) and marine mammals (Ma et al., 2016; Juhel et al., 2021) and to this we now add airborne eDNA detection of elusive bat species.

The use of airborne eDNA also allows for a longer sampling time than visual surveys of bat roosts. During visual surveys, usually the longer a researcher is in the roost the more likely they are to identify all the species present; however, the longer researchers stay inside a roost the more they may disturb the bats, and more manpower is needed to cover more roosts. With airborne eDNA one can leave a sampler in a roost for up to 24 hours (or longer depending on the sampler type and battery) with minimal disturbance to the bats. The units used here emitted no obvious ultrasound (M. Kalcounis-Rueppell pers. Comm) and are quiet at other frequencies; we observed bats roosting directly above them in multiple instances, suggesting they are very minimally disruptive to roosting bats. Such considerations make eDNA samplers ideal for a very non-invasive survey approach. Klepke et al., (2022) found that airborne eDNA accumulates over time, suggesting that the longer a sampler is left in a roost, the more likely it is to capture the total diversity in said roost. Being able to leave samplers in roosts also broadens the potential for simultaneous roost surveys, and the low cost of this sampler design (Garrett et al., 2022) means many can be deployed simultaneously at low cost and risk. This is an inexpensive data-collection method on the ground, allowing a small team to survey many potential roosts simultaneously. Doing so would confirm the species present in both known and suspected roosts, and simultaneously provide preliminary occupancy estimates. This could be especially useful for broad surveys in the neotropics where bat roosts may be hard to find (Villalobos-Chaves et al., 2016).

### Sample Coverage

The number of samplers deployed in the Schoolhouse Cave was sufficient to capture the bat diversity in the roost but did not capture total species richness for all taxa in the area. Many of the other vertebrate species detected in this cave were found presumably as a result of eDNA from the surrounding area drifting into the cave and naturally settling since this cave occupies a physical low point in the natural landscape. It is likely that more sampling is needed to capture the richness of the surrounding area, if not the cave itself. But it is an interesting observation that this low point cave might be a natural eDNA accumulation point if such material drifts and settles out of the air and can become captured in these natural structures. The cow in the attic roost at Helen’s House or the pigs in the caves are false positives for the roost but not for the surrounding area, indicating that eDNA from the surrounding area may accumulate in these locations, making them a better target than open “wind swept” areas. While in most cases we know from alternative data source that our bat detections are consistent with known habitation, our data also suggests that detection should not immediately be used to conclude occupancy and we cannot exclude drift from the local area. This could be true for the detection of *S. parvidens* in the cave-like roosts where other capture methods have failed to indicate such a roosting behaviour in this area. Research investigating how localized airborne eDNA signals are – and how eDNA may move through the environment on wind currents, etc. – will help address such questions and aid in study design.

### Day vs. Night Detections

Patterns of non-uniform DNA shedding have been observed in aquatic eDNA studies (Klymus et al., 2015; Sassoubre et al., 2016; Thalinger et al., 2021) and we suspect a similar pattern here. Bats are more active at night, which may increase eDNA shedding during that time, and thus detection rates may be greater at night. The opposite may be true for farm animals that are diurnal. While our data is based on a single roost (Schoolhouse Cave) over a single 24 hour period, where we had paired day and night measures, we observed a distinct pattern of a greater number of bat detections at night and slightly more non-bat detections in samples collected during the day. We treat this observation with caution, however; while the trend is interesting, the difference observed was not significant. We would not normally report and discuss this non-significant finding but include it here as it may be an important consideration in future sampling designs. Our comparisons are based on six day and six night filters which may not be independent (sampling encompassed only one actual day with three air samplers deployed at the same time) but the emerging pattern is cause for careful consideration of how and when sampling should be conducted. The patterns we observed may indicate that eDNA is a very short-term signal in air, either because of degradation or because it falls out of the air quickly. This has been indicated by the detection of eDNA on leaves (Valentin et al., 2020) where it may be settling out of the air rapidly. It is possible that eDNA signal in air is of short duration and distance and therefore might provide an accurate indication of recent activity. This contrasts with the potential of long-range drift we suspect from the cow eDNA that we detected in some samples. The matrix surrounding these areas of secondary forests includes fields with high cow biomass, a significant and unusual source for DNA in the landscape. Clearly more research is needed to evaluate the role of drift in studies of airborne eDNA.

## Conclusions

We have demonstrated that airborne eDNA can be used to detect vertebrates both inside bat roosts and from the areas surrounding roosts, indicating that airborne eDNA is a potentially game-changing tool for non-invasive surveys of caves, hollows, and roosts. However, more research is needed to understand the ecology of airborne eDNA, including how much eDNA may be drifting into roosts from the surrounding areas, and to determine the best sampling strategy for roost surveys, particularly with respect to sampling intensity, duration, and timing. Our study showcases airborne eDNA as a roost survey tool that could be especially useful in surveying difficult to access locations and determining roost occupancy over periods beyond that of a single visual inventory or camera trapping campaign.

## Acknowledgements

We wish to thank the staff at Lamanai Field Research Center for all their assistance with sampling logistics and research permits. We also thank colleagues who helped with field work and bat captures during the 2022 field season. Thanks to Will Clare, Matt Clare, Kaya Courie, Annie Floyd and Owen Floyd who built and tested the three prototype filters, and Jerry J Grech for the design of our 3D printing. Thanks to Brock and Sherri Fenton for photographs, Helen Haines for access to her attic, and Alejandro Maeda-Obregon for help with the DADA2 pipeline. This work was funded by the Natural Sciences and Engineering Research Council of Canada through the Discovery Grants Program to ELC and an NSERC of Canada CGS-M, Academic Excellence Fund award, York Graduate Scholarship and the Vernon Oliver Stong Graduate Scholarship in Science to NRG.

