## Appendix 1 for "Out of thin air: surveying tropical bat roosts through air sampling of eDNA"

| Species | Sampler A | | Sampler B | | Sampler C | | Sampler D | | Sampler E | | Sampler F | |
|---|---|---|---|---|---|---|---|---|---|---|---|---|
| *Carollia perspicillata* | 0 | 10 | 0 | 12 | 0 | 0 | 0 | 89 | 0 | 0 | 0 | 0 |
| *Glossophaga mutica* | 25847 | 79 | 95389 | 26797 | 49 | 0 | 6508 | 75227 | 8 | 16713 | 21574 | 9989 |
| *Natalus mexicanus* | 0 | 2 | 3672 | 82134 | 39 | 11200 | 17 | 1177 | 66429 | 7241 | 28605 | 10556 |
| *Saccopteryx bilineata* | 12 | 0 | 0 | 0 | 0 | 38 | 0 | 0 | 0 | 0 | 0 | 0 |
| *Sturnira parvidens* | 0 | 0 | 0 | 2265 | 29 | 383 | 0 | 6208 | 0 | 0 | 19860 | 0 |
| *Trachops cirrhosus* | 14 | 0 | 1964 | 0 | 0 | 0 | 14142 | 0 | 0 | 31791 | 0 | 0 |
| *Alouatta palliata* | 0 | 0 | 0 | 20390 | 0 | 0 | 0 | 0 | 0 | 0 | 0 | 0 |
| *Bos taurus* | 35 | 51 | 0 | 16 | 144 | 78 | 0 | 2 | 30865 | 58 | 41073 | 2 |
| *Canis spp.* | 13164 | 65 | 3149 | 699 | 324 | 104327 | 0 | 3006 | 15 | 2395 | 13 | 5625 |
| *Equus caballus* | 0 | 0 | 0 | 0 | 0 | 0 | 4100 | 0 | 0 | 0 | 0 | 0 |
| *Leptodactylus fragilis* | 0 | 0 | 13 | 0 | 0 | 187 | 0 | 0 | 0 | 5255 | 0 | 0 |
| *Ototylomys phyllotis* | 0 | 0 | 0 | 0 | 0 | 0 | 5 | 0 | 13282 | 0 | 23 | 0 |
| *Ovis aries* | 36565 | 127 | 8254 | 0 | 159 | 0 | 49444 | 0 | 6 | 0 | 9 | 0 |
| *Scinax staufferi* | 0 | 0 | 0 | 0 | 0 | 0 | 0 | 0 | 1516 | 0 | 0 | 0 |
| *Sus scrofa* | 55170 | 0 | 0 | 0 | 14 | 112 | 0 | 0 | 0 | 36 | 0 | 0 |
| *Sylvilagus floridanus* | 8820 | 0 | 0 | 0 | 17 | 0 | 0 | 0 | 0 | 0 | 0 | 0 |
| *Trachycephalus typhonius* | 0 | 0 | 3952 | 0 | 0 | 0 | 0 | 0 | 0 | 0 | 0 | 0 |

☐ Day ☐ Night

Figure 1S. The total read count by sampler for each species detected during the day (yellow) and at night (blue) the Schoolhouse Cave.

A)

| Species | Sampler A | | | | | | Sampler B | | | | | | Sampler C | | | | | |
|---|---|---|---|---|---|---|---|---|---|---|---|---|---|---|---|---|---|---|
| | *R1* | *R2* | *R3* | *R1* | *R2* | *R3* | *R1* | *R2* | *R3* | *R1* | *R2* | *R3* | *R1* | *R2* | *R3* | *R1* | *R2* | *R3* |
| Carollia perspicillata | 4544 | 4 | 0 | 0 | 0 | 0 | 0 | 0 | 0 | 7 | 7 | 5 | 0 | 10 | 0 | 8193 | 5 | 0 |
| Glossophaga mutica | 25810 | 22 | 15 | 17 | 32 | 0 | 15082 | 51501 | 28806 | 33 | 10693 | 16071 | 26 | 53 | 0 | 0 | 0 | 0 |
| Natalus mexicanus | 0 | 0 | 0 | 20 | 19 | 0 | 0 | 3672 | 0 | 24381 | 39784 | 17969 | 0 | 2 | 0 | 705 | 10495 | 0 |
| Saccopteryx bilineata | 2 | 0 | 10 | 0 | 0 | 0 | 0 | 0 | 0 | 0 | 0 | 0 | 0 | 0 | 0 | 24 | 0 | 14 |
| Sturnira parvidens | 0 | 0 | 0 | 29 | 0 | 0 | 0 | 0 | 0 | 2265 | 0 | 0 | 0 | 0 | 0 | 0 | 383 | 0 |
| Trachops cirrhosus | 0 | 9 | 5 | 0 | 0 | 0 | 0 | 1964 | 0 | 0 | 0 | 0 | 0 | 0 | 0 | 0 | 0 | 0 |
| Alouatta palliata | 0 | 0 | 0 | 0 | 0 | 0 | 0 | 0 | 0 | 20390 | 0 | 0 | 0 | 0 | 0 | 0 | 0 | 0 |
| Bos taurus | 0 | 16 | 19 | 47 | 97 | 0 | 0 | 0 | 0 | 16 | 0 | 0 | 22 | 29 | 0 | 66 | 0 | 12 |
| Canis spp. | 0 | 0 | 13164 | 85 | 239 | 0 | 0 | 0 | 3149 | 699 | 0 | 0 | 26 | 39 | 0 | 41744 | 8233 | 54350 |
| Equus caballus | 0 | 0 | 0 | 0 | 0 | 0 | 0 | 0 | 0 | 0 | 0 | 0 | 0 | 0 | 0 | 0 | 0 | 0 |
| Leptodactylus fragilis | 0 | 0 | 0 | 0 | 0 | 0 | 0 | 0 | 13 | 0 | 0 | 0 | 0 | 0 | 0 | 0 | 187 | 0 |
| Ototylomys phyllotis | 0 | 0 | 0 | 0 | 0 | 0 | 0 | 0 | 0 | 0 | 0 | 0 | 0 | 0 | 0 | 0 | 0 | 0 |
| Ovis aries | 21229 | 6 | 15330 | 58 | 101 | 0 | 0 | 0 | 8254 | 0 | 0 | 0 | 55 | 72 | 0 | 0 | 0 | 0 |
| Scinax staufferi | 0 | 0 | 0 | 0 | 0 | 0 | 0 | 0 | 0 | 0 | 0 | 0 | 0 | 0 | 0 | 0 | 0 | 0 |
| Sus scrofa | 0 | 55170 | 0 | 0 | 14 | 0 | 0 | 0 | 0 | 0 | 0 | 0 | 0 | 0 | 0 | 0 | 112 | 0 |
| Sylvilagus floridanus | 0 | 0 | 8820 | 17 | 0 | 0 | 0 | 0 | 0 | 0 | 0 | 0 | 0 | 0 | 0 | 0 | 0 | 0 |
| Trachycephalus typhonius | 0 | 0 | 0 | 0 | 0 | 0 | 0 | 3952 | 0 | 0 | 0 | 0 | 0 | 0 | 0 | 0 | 0 | 0 |

Day Night

**B)**

| Species | Sampler D | | | | | | Sampler F | | | | | | Sampler E | | | | | |
|---|---|---|---|---|---|---|---|---|---|---|---|---|---|---|---|---|---|---|
| | *R1* | *R2* | *R3* | *R1* | *R2* | *R3* | *R1* | *R2* | *R3* | *R1* | *R2* | *R3* | *R1* | *R2* | *R3* | *R1* | *R2* | *R3* |
| Carollia perspicillata | 0 | 25 | 18 | 7938 | 17 | 8 | 0 | 0 | 0 | 0 | 0 | 2462 | 0 | 0 | 0 | 0 | 0 | 0 |
| Glossophaga mutica | 0 | 6 | 6502 | 29284 | 39742 | 6201 | 4 | 4 | 0 | 16696 | 8 | 9 | 8 | 21561 | 5 | 9936 | 31 | 22 |
| Natalus mexicanus | 8 | 9 | 0 | 1171 | 6 | 0 | 45759 | 20670 | 0 | 0 | 7241 | 0 | 47 | 28553 | 5 | 7054 | 3491 | 11 |
| Saccopteryx bilineata | 0 | 0 | 0 | 0 | 0 | 0 | 0 | 0 | 0 | 0 | 0 | 0 | 0 | 0 | 0 | 0 | 0 | 0 |
| Sturnira parvidens | 0 | 0 | 0 | 6208 | 0 | 0 | 0 | 0 | 0 | 0 | 0 | 0 | 19860 | 0 | 0 | 0 | 0 | 0 |
| Trachops cirrhosus | 14142 | 0 | 0 | 0 | 0 | 0 | 0 | 0 | 0 | 15446 | 16462 | 0 | 0 | 0 | 0 | 0 | 0 | 0 |
| Alouatta palliata | 0 | 0 | 0 | 0 | 0 | 0 | 0 | 0 | 0 | 0 | 0 | 0 | 0 | 0 | 0 | 0 | 0 | 0 |
| Bos taurus | 0 | 0 | 0 | 2 | 0 | 0 | 0 | 30846 | 19 | 43 | 0 | 15 | 0 | 15 | 41060 | 0 | 0 | 0 |
| Canis spp. | 0 | 0 | 0 | 0 | 0 | 3006 | 0 | 15 | 0 | 0 | 0 | 2395 | 0 | 13 | 0 | 0 | 0 | 5625 |
| Equus caballus | 0 | 0 | 4100 | 0 | 0 | 0 | 0 | 0 | 0 | 0 | 0 | 0 | 0 | 0 | 0 | 0 | 0 | 0 |
| Leptodactylus fragilis | 0 | 0 | 0 | 0 | 0 | 0 | 0 | 0 | 0 | 4181 | 0 | 1074 | 0 | 0 | 0 | 0 | 0 | 0 |
| Ototylomys phyllotis | 0 | 0 | 5 | 0 | 0 | 0 | 0 | 4 | 13278 | 0 | 0 | 0 | 14 | 9 | 0 | 0 | 0 | 0 |
| Ovis aries | 27461 | 21977 | 6 | 0 | 0 | 0 | 6 | 0 | 0 | 0 | 0 | 0 | 9 | 0 | 0 | 0 | 0 | 0 |
| Scinax staufferi | 0 | 0 | 0 | 0 | 0 | 0 | 1516 | 0 | 0 | 0 | 0 | 0 | 0 | 0 | 0 | 0 | 0 | 0 |
| Sus scrofa | 0 | 0 | 0 | 0 | 0 | 0 | 0 | 0 | 0 | 21 | 10 | 5 | 0 | 0 | 0 | 0 | 0 | 0 |
| Sylvilagus floridanus | 0 | 0 | 0 | 0 | 0 | 0 | 0 | 0 | 0 | 0 | 0 | 0 | 0 | 0 | 0 | 0 | 0 | 0 |
| Trachycephalus typhonius | 0 | 0 | 0 | 0 | 0 | 0 | 0 | 0 | 0 | 0 | 0 | 0 | 0 | 0 | 0 | 0 | 0 | 0 |

Day Night

Figure 2S. The total read count by sampler (**A)** samplers A-C, **B)** samplers D-E) and PCR replicate (R#) for each species detected during the day (yellow) and at night (blue) the Schoolhouse Cave.
